# Feature-tuned synaptic inputs to somatostatin interneurons drive context-dependent processing

**DOI:** 10.1101/2025.04.16.649203

**Authors:** William D. Hendricks, Masato Sadahiro, Dan Mossing, Julia Veit, Hillel Adesnik

**Author notes:** These authors contributed equally to this work (William D. Hendricks, Masato Sadahiro).

## Abstract

Mapping neural computation onto the functional microarchitecture of sensory circuits is essential for understanding how brain circuits transform input signals into coherent percepts. Many higher-order perceptual processes emerge in the cortex, yet relatively little is known about how specific connectivity motifs give rise to these computations. To address this challenge, we combined single cell and population-level physiological recordings and perturbation methods to map a context-dependent cortical computation onto the synaptic microarchitecture of the mouse primary visual cortex (V1). We demonstrate a precise, recurrent circuit between cortical pyramidal cells (PCs) and somatostatin (SST) inhibitory interneurons that mediates context-driven figure/ground modulation in V1. Through a like-to-like connectivity rule from PCs and SSTs, this circuit explains SSTs’ visual encoding properties and their resulting impact on contextual modulation in V1. These findings reveal key synaptic and circuit mechanisms that may underlie the earliest stages of scene segmentation in the visual cortex. Moreover, they raise the notion that feature-specific excitatory-inhibitory microcircuitry from PC to SSTs could be a general strategy that the cortex exploits to give rise to higher level computations.

## Introduction

Connectivity patterns between cortical neurons are the building blocks of cortical computations. Yet mapping specific computations onto the synaptic and circuit architecture of cortical circuits has proved challenging. Powerful approaches that correlate physiological response properties of cortical neurons with their connectivity patterns have revealed feature-specific wiring among nearby excitatory neurons ^1–10^, providing a circuit basis for amplification of correlated neural activity. However, these findings have yet to explain higher level computations that may rely on specific connectivity patterns between excitatory neurons and local inhibitory interneurons ^9,11–15^. Moreover, precise perturbations of neural activity are necessary to determine which connectivity features causally drive distinct aspects of cortical computation.

One of the first computations the visual system must perform is segregating figures from the background. A neural correlate of figure segmentation appears as early as primary visual cortex (V1): the response of V1 neurons is much greater to a stimulus within their receptive fields when it constitutes a “figure” than when that same stimulus forms part of the background, a phenomenon known as “figure/ground modulation” ^11,16–19^. Physiologically, a clear neural correlate of this modulation manifests as “orientation-dependent surround suppression” ^17,20–25^. In primates, cats, and mice, studies show that the relative orientation of the center and surround of a visual grating profoundly modulates V1 neural responses ^11,12,16,17,20–22,24–27^. When the center orientation differs from the surround (“cross-oriented”), leading to the percept of a figure, V1 neurons are potently driven ^20–24^. Conversely, when the center orientation matches that of the surround (“iso-oriented”), removing the percept of a figure, V1 neurons are strongly suppressed ^25–28^. Thus, orientation-dependent surround suppression is a widely conserved feature of visual cortical activity that may represent one of the earliest cortical computations supporting the segmentation of the visual scene ^16,19,21,23,24,29^.

Surround suppression in L2/3 depends on the activity of somatostatin-positive GABAergic interneurons (SSTs) ^14,15^. When a grating extends beyond the classical receptive field (CRF) of L2/3 pyramidal cells (PCs), SSTs are potently recruited ^12,14,30,31^ and inhibit both PCs and parvalbumin (PV)-positive interneurons (PVs) ^32–36^. The net inhibition of the PC/PV network suppresses total synaptic activity ^37–39^ in the network which results in greatly reduced spiking for both PCs and PVs. However, when the grating covering the surround does not match that of the center, the spiking activity of PCs and PVs is not suppressed and can even be facilitated ^11,12,20,39^. This orientation-dependent component of surround suppression still lacks a complete synaptic and circuit mechanistic understanding. More generally, we lack a circuit level understanding that explains how figure/ground modulation emerges from the functional microarchitecture of V1 between PCs, SSTs, and other cell types.

Recent studies ^11,12^ have shown that non-matching surrounds drive vasoactive-intestinal peptide (VIP) neurons, which selectively inhibit SSTs ^33,35,40–42^, potentially supporting a model where VIP→SST inhibition is the key means for generating figure/ground modulation. However, optogenetic silencing of VIP neurons only partially reduces the orientation dependence of surround suppression ^11,12^. Thus, whether the inhibitory action of VIP neurons onto SSTs is essential to tune the orientation dependence of surround suppression or whether they play a more modulatory role remains uncertain. An alternative or perhaps even complementary possibility is that SSTs are selectively driven by matching surrounds because only matching surrounds drive sufficiently strong afferent synaptic excitation. This latter model focuses on the excitatory input patterns to SSTs and would likely require a high degree of specificity in their synaptic input architecture: SSTs would have to receive excitation preferentially from pools of PCs with the same orientation preference. Whether this is true is entirely unknown, and so far there is no evidence for such a high degree of specificity of excitatory input to SSTs neurons, or cortical GABAergic neurons in general.

To determine the neural mechanisms of figure/ground modulation we combined extracellular electrophysiology, two-photon calcium imaging, in vivo whole-cell recording, and one and two-photon (2p) optogenetics to map a context-dependent computation onto the synaptic and circuit architecture of mouse V1. We reveal a previously unknown level of specificity in the excitatory input connectivity to SSTs and demonstrate how this circuitry, combined with other recurrent circuits in V1, implements this fundamental visual computation. These findings illuminate a synaptic mechanism for the encoding of contextual information and suggest that such feature-dependent connectivity from PC→SSTs may represent a general principle by which cortical circuits support higher-level computations.

## Results

### Synaptic basis of figure/ground modulation in V1

First, we probed figure/ground modulation in V1 using extracellular electrophysiology. When the orientation of the surround matched that of the center (“iso”), V1 neuron activity was strongly suppressed **(Figure 1A-1C)**. In contrast, when the orientation of the surround was orthogonal to that of the center (“cross”) this suppression largely disappeared. Although this phenomenon is well documented at the spike rate level, its synaptic basis remains poorly understood. Prior work has shown that iso-oriented surrounds drive suppression by decreasing total synaptic input and by lowering the ratio between visually evoked excitation and inhibition ^37^. Cross-oriented surrounds might counteract this suppression by selectively increasing excitatory input, by reducing inhibitory input, or by preventing the surround-induced suppression of total synaptic input. To directly distinguish between these possibilities we made *in vivo* whole-cell recordings from layer 2/3 pyramidal cells (PCs) in awake mice, enabling direct measurement of excitatory and inhibitory postsynaptic currents (EPSCs and IPSCs, respectively) underlying surround suppression. Center gratings evoked strong EPSCs and IPSCs in PCs, while the addition of the iso-oriented surround substantially suppressed both synaptic excitation and inhibition, indicative of network suppression **(Figures 1D and 1E)**. Conversely, addition of a cross-oriented surround restored excitation and inhibition to values indistinguishable to that of the center alone. The results of these intracellular recordings show that cross-oriented gratings avoid driving surround suppression by preventing the network suppression of total synaptic input. Exactly how this occurs, however, requires further investigation into the cell types and cell-type specific connectivity in V1.

**Figure 1:**
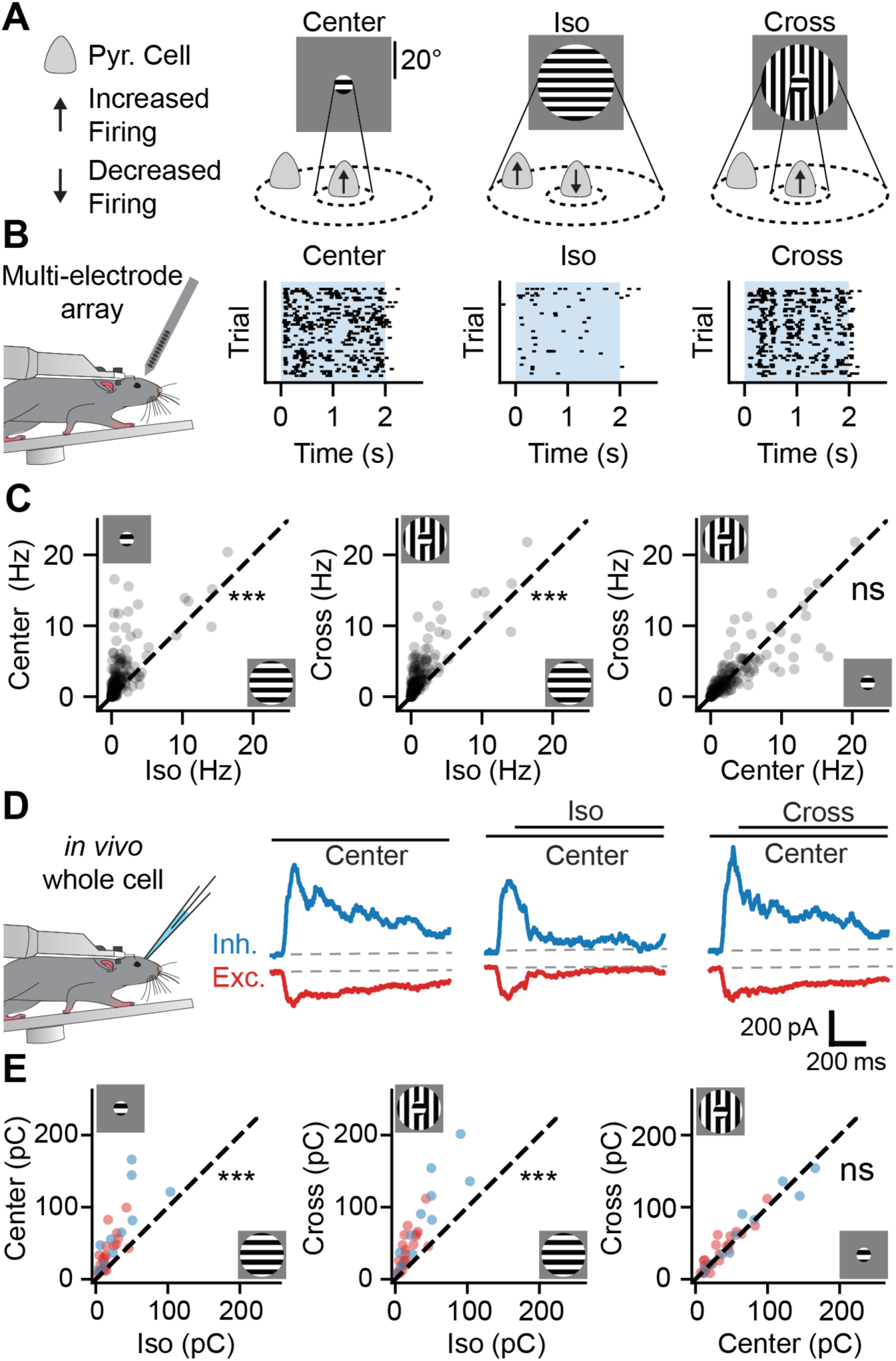
Synaptic mechanism of orientation-dependent surround suppression in the primary visual cortex. **A:** Figure-ground segregation in mouse primary visual cortex (V1) exemplified as orientation-dependent surround suppression. A “center” or “figure” visual stimulus (left) potently drives neural activity, whereas the same stimulus extending into the surround (“iso”, middle) suppresses it. When the surround does not match the center (”cross”, right) neural activity is no longer suppressed. **B:** Extracellular recordings in mouse V1 (schematic, left) in response to “center”, “iso” and “cross” gratings. Raster plots (right 3 panels) are taken from an example regular-spiking (RS) unit. Shaded blue region marks the timing of the visual stimulus. **C:** Scatter plots comparing RS unit responses (n=161 units) to center vs. iso (left, p<10^-5^), iso vs. cross (middle, p<10^-5^), and cross vs. center (right, p=0.676) gratings. P-values were calculated using Wilcoxon signed-rank test and corrected for multiple comparisons using Holm’s method. **D:** Whole-cell voltage clamp recordings from putative pyramidal cells (PCs) in V1 (schematic, left) taken from awake, head-fixed animals. Excitatory (red, n=21 cells) and inhibitory (blue, n=12 cells) current responses to center only, center with iso, and center with cross grating. Note that in these experiments the surround stimulus switched on ∼150 ms after the center. Dashed line represents the baseline holding current for excitatory (-70 mV) and inhibitory (0 mV) potentials. The traces represent the grand average visual responses across cells. Solid black lines above the traces represent timing of the visual stimuli. **E:** Scatter plots comparing synaptic charge-transfer (in pC) of EPSCs (red, n = 21 cells) and IPSCs (blue, n=12 cells) to center vs. iso (left, p<10^-4^), iso vs. cross (middle, p<10^-5^), and cross vs. center (right, p=0.111) gratings. P-values were calculated using Wilcoxon signed-rank test and corrected for multiple comparisons using Holm’s method. ***p<0.001; ns, not significant

### Cortical SST neurons are critical for figure/ground modulation

SST cell activity is required for surround suppression ^14^, but whether they contribute to orientation-tuning of surround suppression is unclear. Thus, we considered the V1 microcircuit and inputs to SSTs **(Figure 2A)** and employed somatic 2-photon calcium imaging in transgenic mice (see Methods). First, we confirmed PCs preferred cross-oriented gratings **(Figure 2B)**, similar to that of our extracellular and intracellular recordings **(Figure 1)**. Notably, however, SSTs strongly preferred iso-oriented compared to cross-oriented gratings **(Figure 2C)**, strikingly opposite to the preferences of PCs **(Figure 2B)**, as well as PVs **(Figure S1)**.

**Figure 2:**
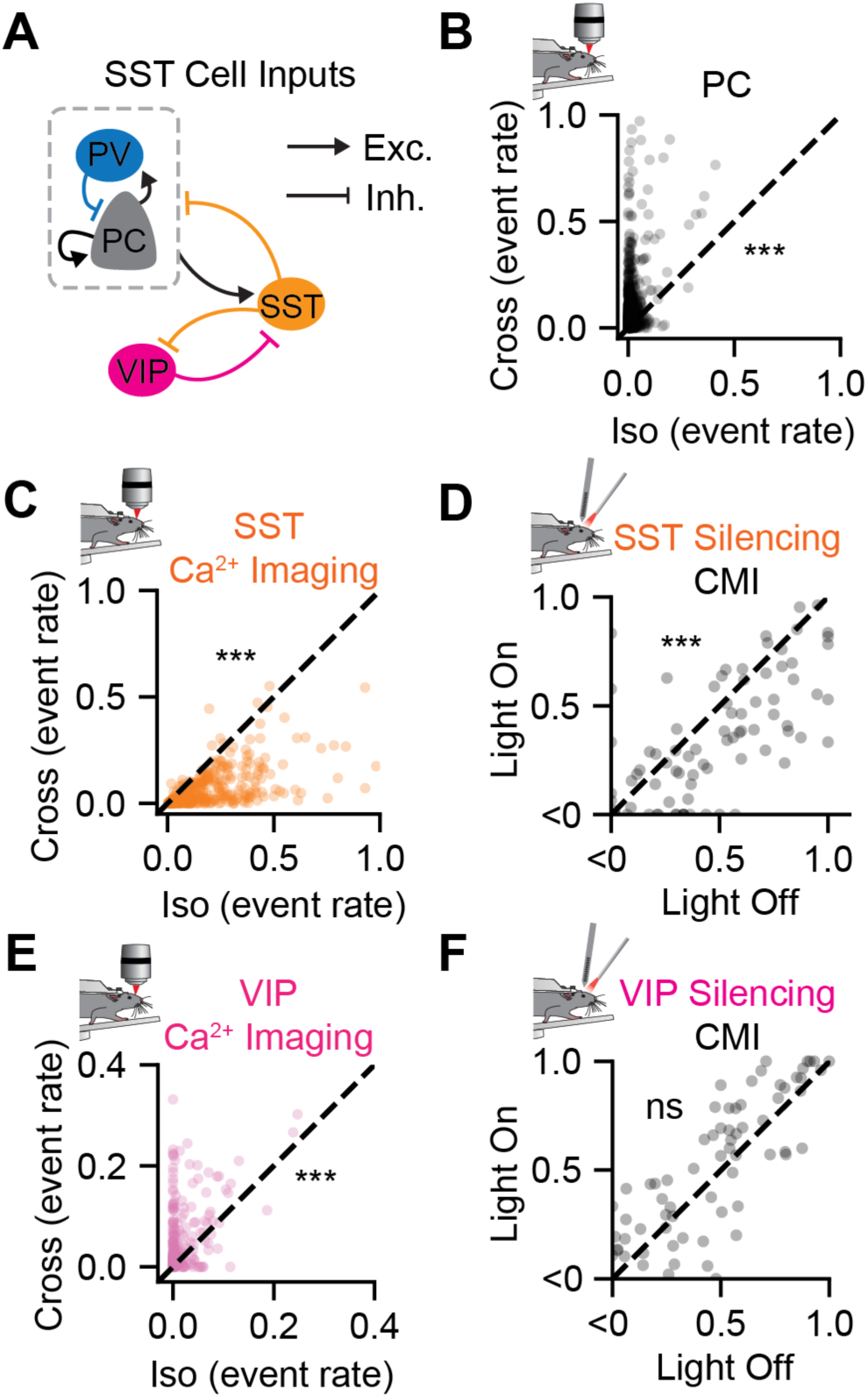
Somatostatin interneurons are required for orientation-dependent surround suppression. **A:** Schematic of the L2/3 V1 microcircuit between PCs (grey), PVs (blue), SSTs (orange), and VIPs (pink). **B:** Scatter plots comparing PC visual responses in the “iso” and “cross” conditions. Activity is measured as the deconvolved event rate, see Methods for details (n=2,329 cells, p<10^-5^, Wilcoxon signed-rank test). Somatic calcium imaging recordings were made from CaMK2a-tTa;tetO-GCaMP6s mice. **C:** Similar to **B**, except for SST cells (n=362 cells, p<10^-5^, Wilcoxon signed-rank test). Recordings were made from SST-IRES-Cre;TITL-GCaMP6s (Ai162) mice. **D:** Scatter plot comparing contextual modulation index (CMI) in the “light on” and “light off” optogenetic silencing conditions of SST cells (n=83 RS units, p<10^-5^, Wilcoxon signed-rank test). Multielectrode array recordings were made from SST-IRES-Cre mice injected with DIO-eNpHR3.0-YFP. **E:** Similar to **C**, for VIP cells (n=318 cells, p<10^-5^, Wilcoxon signed-rank test). Somatic calcium imaging recordings made from VIP-IRES-Cre x TITL-GCaMP6s (Ai162) mice. **F:** Same as in **D**, for VIP silencing (n=71 RS units, p=0.066, Wilcoxon signed-rank test). Multielectrode array recordings were made from VIP-IRES-Cre mice injected with DIO-eNpHR3.0-YFP. ***p<0.001; ns, not significant

To determine whether SST activity is a necessary component of orientation-dependent surround suppression, we used multielectrode arrays to record activity from regular-spiking (RS) putative excitatory neurons and fast-spiking (FS) putative PV interneurons while optogenetically silencing SSTs using eNpHR3.0 (see Methods). If their activity is critical, then silencing SSTs should attenuate orientation-dependent surround suppression. We computed the “contextual modulation index” (CMI) to use as a direct measure of figure/ground modulation. CMI compares the amount of surround suppression between the iso and cross conditions (CMI = (R_cross_ - R_iso_) / (R_cross_ + R_iso_), where R is the mean response during visual stimulation). Thus, a CMI of -1 represents a complete preference to iso gratings, 0 indicates no preference, and 1 represents complete preference to cross gratings. CMI was broadly distributed and almost always positive **(Figure S2)**, reflective of robust orientation-dependent surround suppression.

We found no difference in CMI between FS or RS units **(Figure S1)** or between running states **(Figure S2)**. Strikingly, silencing SST cells significantly decreased CMI, demonstrating that SST activity critically contributes to the orientation-dependence of surround suppression in V1 **(Figure 2D)**.

### An SST “switch-on” model for figure/ground modulation

Based on the foregoing data, we can conceive of a model in which iso-oriented surround stimuli recruit SST neurons, driving network suppression of total synaptic activity, thereby reducing network firing rates. Cross-oriented stimuli then would not recruit SST neurons, avoiding this network suppression, leaving firing rates intact, and give rise to figure/ground modulation. However, this conceptual model still requires a synaptic mechanism that explains the selective recruitment of SST neurons for iso-oriented but not for cross-oriented stimuli. We can consider two scenarios: first, vasoactive intestinal peptide (VIP) interneurons, which preferentially inhibit SSTs, might specifically “switch-off” SSTs for cross-oriented stimuli thereby preventing surround suppression ^11,12^. Alternatively, iso-oriented gratings, but not cross-oriented gratings, could specifically “switch-on” SSTs and thereby engage surround suppression. In the first case, a surround grating of any orientation is sufficient to excite SSTs, but VIP activity cancels this excitation. In the latter case, only iso-oriented gratings generate sufficient excitation in SSTs to drive firing. Moreover, both of these mechanisms could exist in tandem, operating synergistically with one another to modulate figure/ground perception.

To examine the “switch-off” hypothesis we measured visual responses of VIP neurons to iso- and cross-oriented stimuli and probed the network impact of optogenetically silencing their activity. Two photon calcium imaging from VIP neurons showed that nearly all VIP neurons responded more strongly to cross-oriented compared to iso-oriented stimuli **(Figure 2E)**, consistent with prior findings ^11,12^. To test whether VIP neurons have a causal role in the orientation dependence of surround suppression, we optogenetically suppressed their activity and measured the impact on V1 activity and CMI. Illumination of VIP neurons expressing the silencer eNpHR3.0 strongly suppressed their visually evoked activity and reduced PC activity **(Figure S3)**, confirming the efficacy of our manipulation. Surprisingly, despite the strong effect that VIP silencing had on V1 firing rates in both the cross and iso conditions **(Figure S3)**, our manipulation had no detectable effect on CMI **(Figure 2F)** implying that VIP cell activity is not essential for the orientation-dependence of surround suppression. Given two prior studies found that optogenetically suppressing VIP cells altered this contextual modulation ^11,12^, we considered the possibility that an effect on CMI was not detectable in our sample size. Thus, we executed an analogous experiment optogenetically suppressing VIPs cells while sampling a much larger pool of PCs with somatic 2p calcium imaging as done in one of these prior studies ^12^. In this case, we found VIP silencing drove a statistically significant, albeit partial effect on CMI **(Figure S4)**, both qualitatively and quantitatively consistent with prior work ^11,12^. However, the remaining contextual modulation after VIP suppression suggests there must also be an alternative mechanism that can explain orientation-dependent surround suppression in V1 L2/3 PCs.

Thus, we sought to test the alternative hypothesis that iso-oriented gratings selectively switch-on SSTs by driving substantially more excitation in SSTs than do cross-oriented gratings. We targeted SST neurons for whole-cell recordings under two-photon guidance **(Figure 3A and 3B)** and probed their synaptic excitatory input to small gratings, iso-oriented gratings, and cross-oriented gratings. We presented each stimulus across four orientations and focused on the response pattern at each SST’s preferred orientation. Consistent with SSTs’ well-known spike-rate preference for large compared to small stimuli ^12,14,30,43^, large gratings drove substantially more synaptic excitation than did small gratings **(Figure 3D)**, opposite to what we found in PCs **(Figure 1)**. More importantly, we found that SSTs also received substantially more excitation for iso-compared to cross-oriented gratings **(Figure 3E)**, while cross-oriented gratings drove similar excitation to the center alone **(Figure 3F)**. To test whether excitatory input to SSTs largely originates from the surround, we compared the response to large gratings to the same stimulus but with the center removed (“aperture gratings”). Remarkably, aperture gratings drove nearly as much excitation as iso-oriented gratings **(Figure 3G)**, implying that most of SSTs’ excitatory input derived from the retinotopic surround. These measurements of synaptic excitatory input to SSTs provide a clear explanation for why iso-oriented but not cross-oriented gratings strongly recruit SST activity.

**Figure 3:**
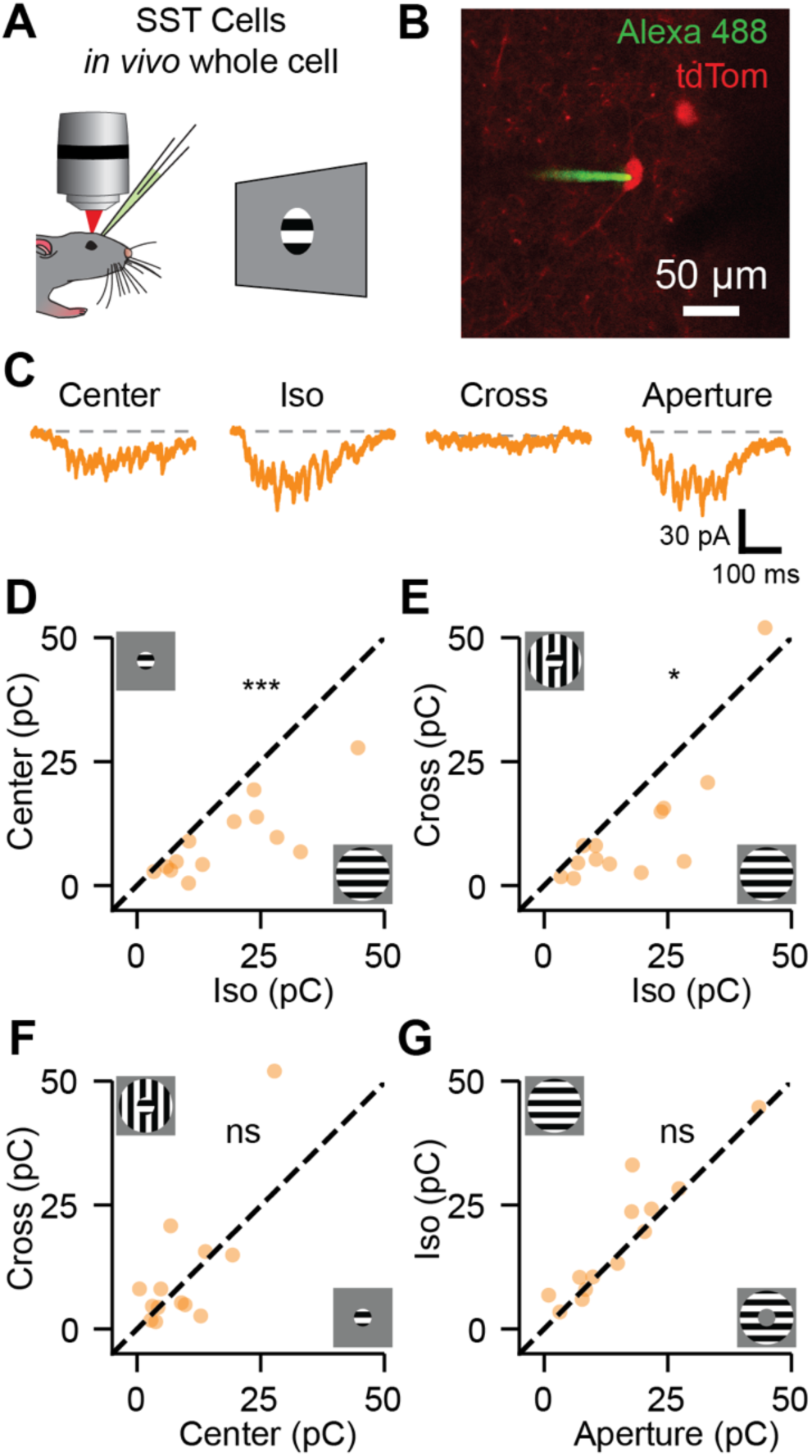
SST cells receive orientation-tuned excitatory input from the retinotopic surround. **A:** Schematic of 2-photon guided *in vivo* whole-cell voltage clamp recording of SST cells in SST-IRES-Cre;Rosa-LSL-tdTomato mice. **B:** Representative image of a dye-filled patch pipette recording a tdTomato+ SST neuron. **C:** Example visually evoked synaptic EPSCs from an SST neuron to the indicated visual stimuli. Dashed gray line represents baseline holding potential (-70 mV). **D:** Scatter plots comparing synaptic charge-transfer (in pC) of EPSCs in SST neurons to center vs. iso stimuli (n=13 cells, p=0.00098, Wilcoxon signed-rank test). **E:** Similar to **D**, but comparing cross vs. iso stimuli (n=13 cells, p=0.018, Wilcoxon signed-rank test). **F:** Similar to **D-E**, but comparing cross vs. center stimuli (n=13 cells, p=0.839, Wilcoxon signed-rank test). **G:** Similar to **D-F**, but comparing cross vs. aperture stimuli (n=13 cells, p=0.188, Wilcoxon signed-rank test). *p<0.05, ***p<0.001; ns, not significant

### Feature-dependent cortical microcircuitry for figure/ground modulation

What features of the synaptic cortical microarchitecture can explain how SSTs’ receive more visually evoked excitation for iso- compared to cross-oriented grating? SSTs receive minimal synaptic input from L4 but substantial excitation from within L2/3 ^14,35,44,45^. If populations of L2/3 PCs with common orientation tuning converge onto individual SSTs, this would explain why they receive more synaptic excitation for iso-oriented gratings than other stimuli. Thus, we hypothesized that lateral input within L2/3 between PCs and SSTs should be ‘like-to-like’ in orientation space; that is, PCs should selectively drive SSTs with common orientation preferences. To test this, we used 2p holographic optogenetics to photo-activate individual orientation-tuned PCs while performing simultaneous 2-photon calcium imaging of PCs and SSTs expressing GCaMP6 **(Figure 4A)**. We conditionally expressed ChroME2s ^46^ in PCs for photostimulation along with blue fluorescent protein (BFP) to positively identify SSTs *post-hoc* **(Figure 4B)**. First, we confirmed the efficacy of photo-stimulating PCs using targeted holographic illumination **(Figure 4C)**. Then, we confirmed that we could robustly drive visual responses in both PCs and SSTs **(Figure 4D)** and then mapped the orientation tuning of PCs and SSTs using full-screen high-contrast drifting gratings **(Figure 4E and 4F)**. Finally, we photo-activated a sequence of individual PCs while monitoring the somatic calcium activity of nearby SSTs **(Figure 4G and 4H)**. Consistent with our “switch-on” hypothesis, the shared orientation preference of PCs and SSTs strongly predicted their functional connectivity: co-tuned PCs drove substantial activation of SSTs, while orthogonally tuned PCs drove slight suppression **(Figure 4I and 4J)**. These results provide the first demonstration of a “like-to-like” circuit motif between cortical excitatory neurons and SST interneurons. Taken together, our data provide a clear mechanism that connects precise patterns of synaptic connectivity in V1 to recurrent circuit dynamics that help mediate higher-order contextual computations.

**Figure 4:**
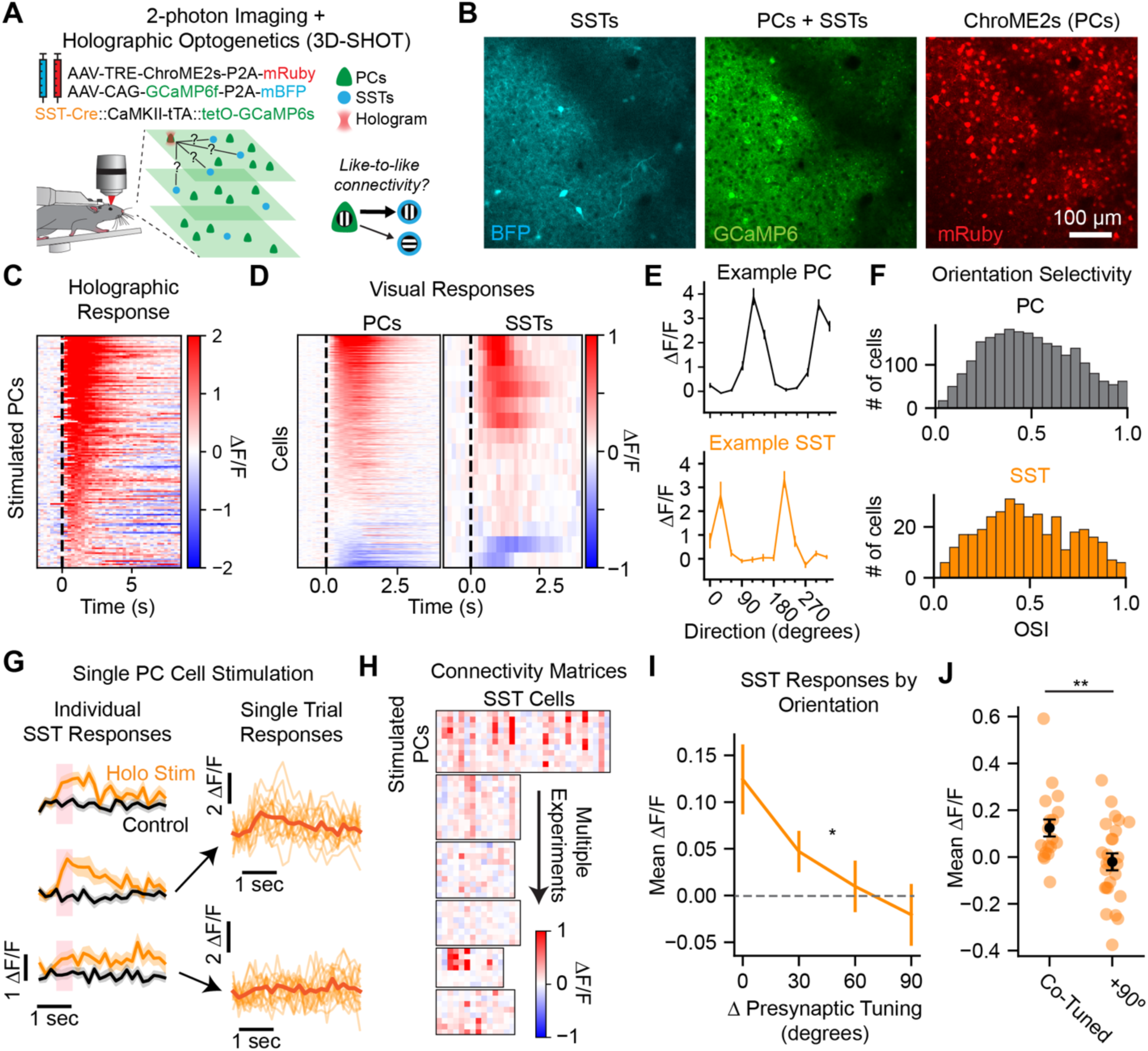
2-photon optogenetic mapping reveals like-to-like connectivity between PCs and SSTs. **A:** Schematic depicting the mapping of functional connectivity between PCs and SSTs with 2-photon holographic optogenetics and calcium imaging. Mice were injected with a viral cocktail that enables Cre-conditional expression of GCaMP6f and BFP in SSTs and TRE-conditional expression ChroME2s opsin in PCs. **B:** Representative images front the same field of view of a mouse expressing BFP in SSTs, GCaMP6 in PCs and SSTs, and ChroME2s (mRuby3) in PCs. Scale bar applies to all images. **C:** Example calcium responses of targeted PCs (n=150 cells) to 2p holographic stimulation. Single neurons were targeted with 50 mW. **D:** Example visual responses of PCs (left, n=368 cells) and SSTs (right, n=16 cells) to high-contrast full-screen drifting gratings. All detected ROIs in the FOV are plotted. **E:** Representative orientation tuning curves from PCs (top, black) and SSTs (bottom, orange). **F:** Histogram of orientation selectivity index (OSI) for PCs (top, grey, n=2,350 cells) and SSTs (bottom, orange, n=362 cells) **G:** Example calcium traces of three SST cells in response to 2p holographic PC stimulation (left column). Single PCs were optogenetically stimulated with 50 mW of light (red shading). Control trials (black traces, no stimulation) were interleaved with holographic stimulation trials (orange traces). Both average traces (left column) and traces from individual trials (right column) are shown. Average traces (bold orange line) are also overlaid on individual trials (right column). **H:** Connectivity matrices from 6 exemplary experimental sessions. Each pixel represents the mean response of an SST (columns) to holographic stimulation of a stimulated PC (rows). Each box comes from an individual connectivity mapping session (n=4 mice shown). **I:** Quantification of SST responses (mean ΔF/F) during 2p holographic stimulation of putative pre-synaptic PCs, sorted by relative difference in orientation (in visual degrees) to stimulated PCs (n=120 total tested PC-SST pairs, 7 sessions, 4 mice, p=0.0247, Kruskal-Wallis H-test). **J:** Similar to **I**, except plotting the individual responses of SSTs to either co-tuned or orthogonally-tuned (Δ +90°) (co-tuned, n=18 PC-SST pairs; +90°, n=26 PC-SST pairs, p=0.00674, Mann-Whitney U-test). *p<0.05, **p<0.01

## Discussion

Our approach for probing the synaptic and circuit logic of figure/ground modulation reveal previously unknown functional principles of cortical wiring and provide a detailed mechanistic model of a cortically emergent computation. Based on the aforementioned experiments, we propose a conceptual model to explain figure/ground modulation: iso-oriented stimuli drive co-tuned PCs across the retinotopic map in the visual cortex, and these co-tuned PCs selectively converge onto individual SSTs, causing them to selectively respond to iso- but not cross-oriented stimuli. In turn, this population of co-tuned SSTs inhibits PCs (as well as PVs and VIPs), suppressing total synaptic activity and network firing rates in the absence of a perceptual figure **(Figure 5A)**. Conversely, cross-oriented stimuli drive orthogonally tuned pools of PCs between the retinotopic center and surround, and as a consequence SSTs receive less overall excitation and fire much less when the image contains a perceptual figure. This prevents network suppression by SSTs and results in stronger overall V1 activity **(Figure 5B)**. These high firing rates presumably help define the existence of specific figural objects in the scene. Inhibition from VIPs onto SSTs, specifically for figure-containing stimuli ^11,12^, may still serve to enhance the consequences of feature-dependent excitatory wiring between PCs and SSTs. The net consequence of the synaptic microarchitecture described here may be one of the first physiological steps in figure/ground modulation.

**Figure 5:**
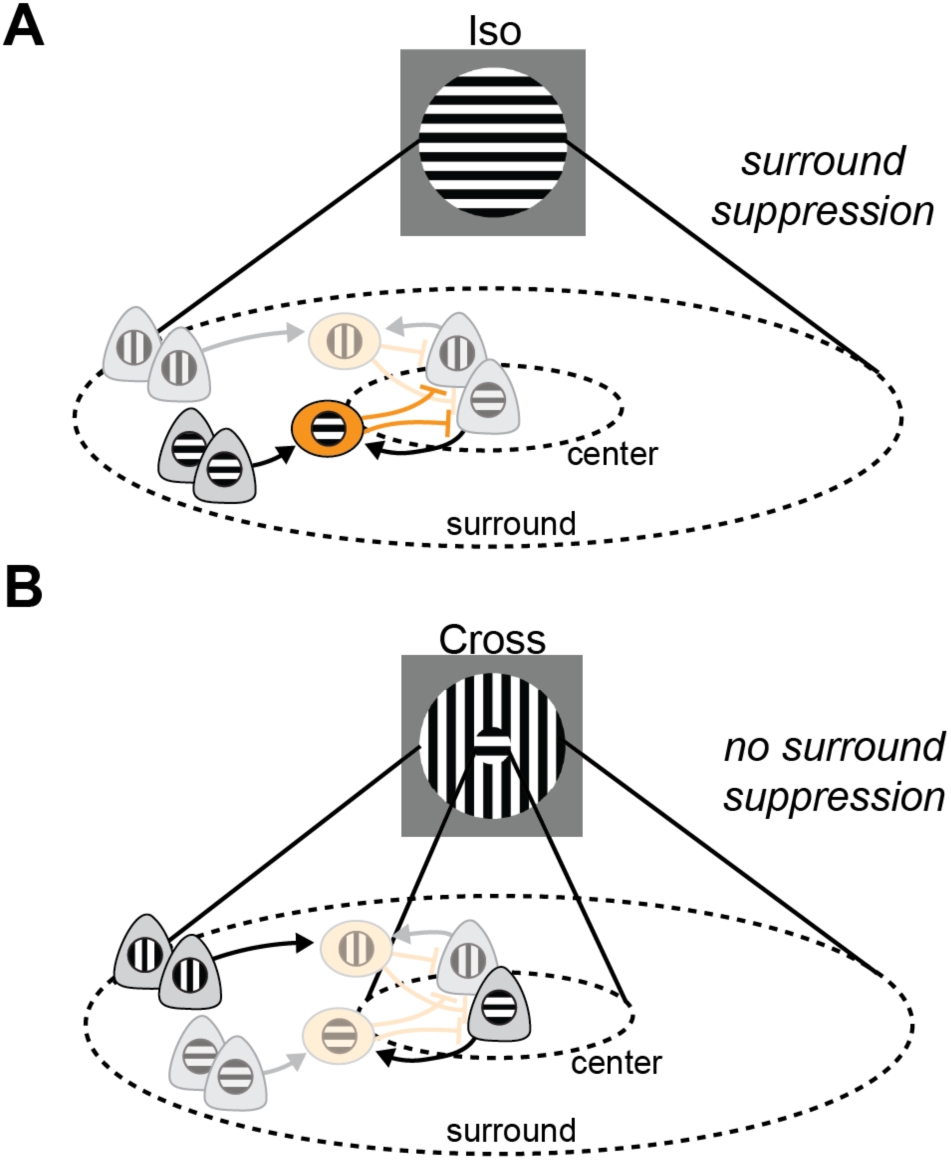
Like-to-like functional connectivity model for orientation-dependent surround suppression. **A:** Tuned (“like-to-like”) PC→SST connections from both the center and surround combine and sufficiently increase SST activity and drive surround suppression to iso gratings. Some connections are omitted for clarity. Shading represents relative activity of connections (arrows) and cells. Light shading represents present, but not active connections and cells. Darker shading represents present and active connections and cells. Center and surround receptive fields are indicated by dashed lines. **B:** For cross-oriented gratings, surround PCs and center PCs drive synaptic input into 2 different populations of SSTs, but not enough to sufficiently increase SST activity for surround suppression.

Since the basic features of cortical architecture are preserved across widely different cortical areas ^47,48^, our work raises the notion that feature-specific lateral wiring from PCs to SSTs might generally be involved in higher order cortical computations.

SSTs also derive substantial input from top-down feedback pathways from higher cortical areas ^49–53^, where excitatory neurons often show more integrative properties ^54^. Suppressing top-down feedback reduces both surround suppression across multiple species and can also reduce its orientation dependence ^11,55–57^. This suggests that excitatory feedback to V1 SSTs might also obey a ‘like-to-like’ rule. So far we lack the tools to map the feature-dependence of functional connectivity between distant excitatory neurons and SSTs, or any other cell type. However, this may be possible with recent advances in very large field of view 2p imaging and 2p optogenetic platforms ^58^, or through a combination of transsynaptic tracing strategies ^6,10,59^ and optogenetic perturbations. Moreover, while the increasingly well-characterized VIP disinhibitory circuit ^12,33,60,61^ is not required for setting up the computation we study here, it likely acts as a potent brain-state or context-dependent modulator that could scale the gain of the feature-selective PC→SST circuit to match ongoing behavioral demands ^30,60,61^.

Moreover, since the cortical neuromodulatory state can control the dynamics of synapses between PCs and SSTs ^62^, it is possible VIPs play a more crucial role when the excitatory synaptic gain onto SSTs changes across different behavioral states.

We note that our conceptual model **(Figure 5)** does not require like-to-like connectivity from SST→PC although it does not exclude it. If SSTs broadly synapse onto PCs without respect to their shared tuning preferences, orientation-dependent surround suppression will still arise if SST firing requires the conjunction of synaptic input from both surround and center PCs that are co-tuned to the SST. In this scenario, different pools of orientation-tuned SSTs fire for different orientations of background iso-stimulus, but for cross-oriented stimulus few if any SSTs are strongly recruited, consistent with our observation that SSTs receive strong synaptic input from the surround **(Figure 3).** Although “like-to-like-to-like” functional connectivity in the PC→SST→PC pathway may not be necessary, it might reinforce orientation-dependent surround suppression that is initially set up by the patterns of excitatory synaptic input to SST neurons we described here. Further experiments mapping SST→PC functional connectivity pattern will help elucidate the circuit motifs that support orientation-dependent surround suppression.

Contextual modulation includes other key computations beyond figure/ground modulation. The visual system must also address sensory ambiguities due to occlusion, low contrast, variable reflectance, etc. to identify objects. Top-down feedback circuits ^52,53,63,64^, together with lateral connections within V1 ^11,12,34,35^, might address all of these challenging sensory contexts through precise patterns of synaptic connectivity between excitatory and specific subtypes of inhibitory interneurons. Other sensory cortical areas, and even higher cortical regions, contain similar sets of cell types with potentially similar patterns of connectivity ^40,47,48,65–67^. Thus it may be possible that cortical areas might repeatedly exploit the types of functional motifs we discovered here to implement a broad array of critical computations and defects in these circuits might help explain features of complex cognitive or neurological disorders.

## Methods

### Animals

All experiments were approved by the University of California, Berkeley Animal Care and Use Committee. All mouse strains were purchased from Jackson labs. Strains used in experiments are: Sst-IRES-Cre (RRID:IMSR_JAX:013044), Vip-IRES-Cre (RRID:IMSR_JAX:010908), VIP-IRES-Cre; SST-IRES-Flp (RRID:IMSR_JAX:031629), Camk2a-tTa (RRID:IMSR_JAX:007004), tetO-GCaMP6s (RRID:IMSR_JAX:024742), Rosa-LSL-tdTomato (RRID:IMSR_JAX:007909), and Ai162 (TIT2L-GC6s-ICL-tTA2) (RRID:IMSR_JAX:031562). Experiments were performed on equal numbers of adult male and female mice.

### Viral injections

SST- and VIP-IRES-Cre mice used for electrophysiology were injected neonatally (P3-P5) with AAV-hSyn-DIO-eNpHR3.0-YFP. For VIP silencing experiments, VIP-IRES-Cre; SST-IRES-Flp mice were injected neonatally with AAV9-CAG-DIO-eNpHR3.0-mRuby3, as previously described ^43,68^ and then as adults with AAV9-hSyn-GCaMP6s (UPenn Vector Core) immediately prior to cranial window placement ^43^. For 2-photon holographic optogenetic experiments, SST-Cre; CaMK2a-tTA;tetO-GCaMP6s mice were injected with a cocktail of AAV-TRE-ChroME2s-H2B-mRuby3 and AAV-hSyn-DIO-GCaMP6f-p2A-mTagBFP2 ^3^. In brief, for adult viral injections, mice 6 weeks or older were anesthetized with 2% isoflurane, given 2 mg/kg of dexamethasone to control cerebral edema and 0.5 mg/kg of buprenorphine as perioperative analgesia, and place in a stereotaxic instrument for head-post implantation, viral injections, craniotomy, and window placement. Body temperature was maintained at 37° C. A small incision was made to remove the scalp and retract the fascia, and 2-3 burr holes (minimum 1 mm apart) were made with a dremel over left V1 (2.7 mm lateral, 1 mm anterior, of lambda). Virus was slowly injected (300 nL, 50 nL/min; Micro4, World Precision Instruments) through each burr hole via a beveled glass pipet (Drummond Scientific).

Headplating and cranial windowing were performed immediately following viral injections. The skull was lightly scored with a drill bit and vetbond applied to improve adhesion of the dental cement and headplate. A 3.5 mm diameter opening in the skull was made with a biopsy punch and a cranial window consisting of two stacked 3 mm diameter coverslips and one 5 mm diameter coverslip on top was secured into place with dental cement (Metabond). Finally, a custom titanium headplate was cemented into place. Mice were given postoperative analgesics (0.5 mg/kg buprenorphine and 5 mg/kg meloxicam) and monitored during recovery.

### Electrophysiology

#### In vivo whole-cell patch clamp recording in V1

*In vivo* awake whole-cell voltage clamp experiments from V1 L2/3 pyramidal cells (Figure 1) in mice were prepared as previously described ^37^, following several days habituation to head restraint on a free spinning running wheel. On the day of recording, mice were briefly anesthetized with 2% isoflurane, the skull over the primary visual cortex was thinned, and a small (< 300 µm) craniotomy was performed with a 27 gauge needle for electrode access. The dura was not removed. Mice were then fixed on the recording set up and allowed to recover for 10-15 minutes after which they ran freely on the circular treadmill. Patch electrodes (4-5 MΩ) were inserted perpendicularly to the cortex and advanced into L2/3 (< 350 µm after first contact with the pia). Cells were targeted with the standard “blind” technique ^37,69^. Pipettes were filled with a cesium based intracellular solution also containing QX-314 and TEA (Sigma-Aldrich) to block active conductances. Excitatory currents were measured at a holding potential of -70 mV, while inhibitory currents were measured at excitatory reversal (∼+10 mV without correction for the junction potential). Access resistance ranges from ∼20-35 MΩ and was partially compensated for via the Axopatch 200B (Molecular Devices) amplifier.

After establishing whole-cell configuration at -70mV holding potential, the visual receptive field (RF) was mapped by presenting a drifting grating stimulus (12 visual degrees) on a gamma corrected 23-inch LCD display (Eizo FORIS FS2333) positioned 10 centimeter from the contralateral eye, and manually moving the stimulus to the location and rotating to the preferred orientation that evoked the maximum excitatory postsynaptic current. Visual stimuli were generated with the Psychophysics Toolbox ^70^.

#### Extracellular multi-unit electrode recording in V1

For extracellular multi-unit electrode recordings **(Figure 1, 2)**, mice were prepared similarly as above. Rather than a patch electrode, one or two 16-channel linear electrodes with 25 micron spacing (NeuroNexus, A1x16-5mm-25-177-A16) were guided into the brain using micromanipulators (Sensapex) and a stereomicroscope (Leica) up to the depth of Layer 4 (450 µm). Electrical activity was amplified and digitized at 30 kHz (Spike Gadgets), and stored on a computer hard drive. The cortical depth of each electrical contact was determined by zeroing the bottom contact to the surface of the brain. Electrodes were inserted close to perpendicular to the brain’s surface for single electrode recordings and ∼25 degrees from vertical for the two electrode experiments. After each recording a laminar probe coated with the lipophilic dye DiI was used to mark each electrode track to quantitatively assess insertion angle and depth with post-hoc histologic reconstructions. The laminar depth of recorded units was corrected for the insertion angle and the local curvature of the neocortex.

Visual stimuli were generated with the Psychophysics Toolbox on an Apple Mac Mini and were presented on a gamma corrected 23-inch LCD display with a 60-Hz refresh rate (Eizo FORIS FS2333). At the beginning of each recording session the receptive fields of MUA recorded at each cortical location were mapped with sparse noise to be able to precisely position the grating stimuli. The stimulus was centered on a location where a small grating, movable by hand, elicited a clear response. Sparse noise consisted of black and white squares (2 visual degrees, 80 ms) on a 20° x 20° grid (in visual degrees) flashed onto a gray background of intermediate luminance. To improve receptive field estimation the same stimulus grid was offset by 1 degree and the resulting maps were averaged. MUA average receptive fields were calculated by reverse correlation. MUA activity was immediately analyzed offline and the grating stimuli were then centered on the peak of the resulting population receptive field.

The center-surround stimulus consisted of full contrast drifting square-wave gratings at 0.04 cycles per degree and 2 cycles per second presented for 2s with at least 1s inter stimulus interval. The center grating was presented at eight (0°-315° in steps of 45° - SOM-Cre population) or four (0° - 270° in steps of 90° - VIP-Cre population) different orientations, with a circular aperture of 8-20° visual degrees diameter, centered on the MUA receptive field, which was surrounded by a 60 degree grating with either 0° or 90° offset (SST-Cre population) or one of seven different relative orientations (0-90° in steps of 15°, VIP-Cre population). We also presented the center grating without surrounding grating (center-only condition).

#### Optogenetic stimulation for extracellular multi-unit electrode recording in V1

For optogenetic stimulation of eNpHR3.0 in vivo (Figure 2) we used red (center wavelength: 625 nm) from the end of a 1-mm diameter multimode optical fiber coupled to a fiber coupled LED (0.39 NA, Thorlabs) controlled by digital outputs (NI PCIe-6353). The fiber was placed as close to the craniotomy as possible (<3 mm). The illumination area was set to illuminate a wide area including all of V1. Light levels were tested in increasing intensities at the beginning of the experiment and were kept at the lowest possible level that still evoked observable change in ongoing activity for the remainder of the recording. Light power at the tip was ∼4-8 mW. The red LED was switched on for 1 s starting 0.5 s after start of the visual stimulus in 50% of the trials. The period of light was chosen to influence the stable steady-state of the response to the grating and all analysis was performed during this time window.

#### In vivo targeted two photon patch clamp recording from V1 SST neurons

In vivo two photon targeted whole-cell voltage clamp experiments from SST neurons **(Figure 3)** were performed in SST-Cre mice crossed with Rosa-LSL-tdTomato (SST-Cre;Ai9). Preparatory surgery of implantation of a head post with an integrated recording chamber and a 3 mm craniotomy were performed under 2% isoflurane anesthesia, sedation under chlorprothixene (5mg/kg; intraperitoneal; Sigma-Aldrich), and cerebral edema reduction with dexamethasone (2 mg/kg; subcutaneous). The dura was removed using fine forceps. The craniotomy was left open but covered and protected with 0.9% low-melting point agarose and hydrated with artificial cerebrospinal fluid. Following surgery, mice were maintained under anesthesia during recording at 0.25-0.5% isoflurane. Mouse body temperature was maintained with a feedback-controlled heating pad. Under two photon visual guidance, a patch electrode (5-6 MΩ) containing cesium-based intracellular solution (containing QX-314 and TEA; Sigma-Aldrich) mixed with AlexaFluor 488 (50 µM; Thermo Fisher Scientific) was directed to the cell along an oblique angle of 24°. Continuous positive pressure was applied to the patch electrode during guidance through brain tissue to ensure visibility and clearance of the electrode tip. Contrasts in fluorescence created by background fluorescence of tdTomato and AlexaFluor 488 were used to avoid blood vessels and identify the soma of non-SST neurons. Sealing attempts were only made on somata with tdTomato expression. Access resistance ranges from ∼20-35 MΩ and was partially compensated for via the Multiclamp 700B amplifier (Molecular Devices).

After establishing whole-cell configuration at -70mV holding potential, the RF of the SST neuron was mapped by presenting a drifting grating stimulus (12 visual degrees), on a Retina iPad LCD display (Adafruit Industries) positioned 10 centimeter from the contralateral eye, and manually moving the stimulus to the location and rotated to the base orientation (0°) that evoked the maximum excitatory postsynaptic current.

Finally, visual evoked excitatory currents in the RF-centered SST neurons were measured in response to different visual stimulus types (“center”, “iso”, “cross”, “aperture”) across 2 orientations, 0° and orthogonal (90°). Visual stimulus center was defined as the stimulus space occupying the central 12 visual degrees, and the surround, present in “iso”, “cross”, and “aperture” was defined as the stimulus space beyond center up to 60 visual degrees. All visual stimuli were generated and presented at 1 Hz, 0.08 cycles per degree, 100% contrast using PsychtoolBox (v3.0).

### One-photon optogenetics

Calcium imaging and one photon optogenetics were performed as previously described ^43^. Briefly, mice were headfixed and allowed to run freely on a rotating circular treadmill under a 16x magnification objective (Nikon) and two-photon resonant scanning microscope (Neurolabware). One-photon illumination (617 nm LED, Thorlabs) for optogenetic silencing was filtered by a 613/22 nm single-bandpass filter (Semrock) and delivered through the objective at 6 mW. The PMT was protected by a shortpass filter and any remaining artifacts were subtracted in post-processing. Visual stimuli were presented using Psychtoolbox 3 ^70^ on a LCD monitor placed 13-15 cm from the eye.

Receptive fields were mapped using 10° drifting grating patches (0.08 cycles per degree, 1 Hz temporal frequency) that rotated in direction 360° over the course of 1.5 seconds. Patches were randomly interleaved within a 40° x 40° grid at 5° intervals.

### Two-photon holographic optogenetics

Calcium imaging and two photon holographic optogenetics were performed similarly as previously described ^3,71^, on a microscope (Sutter MOM, Sutter Instruments) adapted to be capable of simultaneous 3D imaging (resonant-galvo scanning: 3 z-planes, 30 Hz frame rate, 6.36 Hz volume rate) and 3D photostimulation (3D-SHOT) ^3,71–73^. Images were collected using ScanImage 2019 (MBF Biosciences, formerly Vidrio Technologies) with a 20X (1.0 NA) magnification water-immersion objective (Olympus). A dual band-pass dichroic mirror (Chroma, 59003m) was placed prior to the PMTs to direct blue and red light into the same PMT, avoiding GCaMP fluorescence contamination when imaging BFP cells at 830 nm, and thereby enhancing our ability to accurately detect BFP+ neurons. A swapable blue or red single band-pass filter was then used when collecting BFP or mRuby images, respectively. An electrically-tunable lens (Optotune) was placed in the imaging path immediately prior to the resonance-galvo mirrors for fast-z scanning. Imaging laser (Chameleon Ultra II, Coherent) power was restricted to <75 mW at 920 nm to minimize scanning induced crosstalk. All calcium imaging experiments were performed in L2/3 of V1. For experiments with holographic stimulation, custom MATLAB control software was used to control holographic stimulation and synchronize image acquisition.

Holographic optogenetic stimulation was performed using a Monaco 40W laser (Coherent) directed into our custom-built 3D-SHOT optical path and temporally focused using a blazed diffraction grating (Newport Corporation). The beam was directed through a rotating diffuser to randomize the phase and expand the beam onto the spatial light modulator (SLM) (Meadowlark, HSPDM-1K). Phase masks were calculated using a weighted Gerchberg-Saxton algorithm and displayed on the SLM to generate temporally focused spots in 3D. Imaging and photostimulation paths were merged prior to the tube lens with a polarizing beamsplitter (Thorlabs). Photostimulation was synchronized to the scan phase of the resonance mirror and gated with an Arduino Mega to limit stimulation artifacts to the very edge of the imaging FOV. Tiffs were cropped post-hoc to remove any remaining contamination. Imaging and photostimulation paths were first manually co-aligned then digitally calibrated ^3^ to achieve precise targeting and ensure an even distribution of power throughout the FOV. The calibration was verified prior to every experiment by burning holes in a thin fluorescent slide. Digital offsets were applied to the calibration to account for minor drifts (<5 µm) in co-alignment. Larger misalignments triggered a full recalibration. Slow drifts of the FOV over the course of the experiment were manually corrected, aided by the location of blood vessels and other landmarks.

Visual stimuli were presented on a 2,048 x 1536 Retina iPad LCD display (Adafruit Industries) placed 10 cm from the mouse. The display was synchronized with the galvos such that the backlight was only illuminated during the X- (resonant) galvo turnaround time to prevent light from the monitor contaminating the 2p imaging. Drifting gratings (50 visual degrees, 1 Hz, 0.08 cycles per degree, 100% contrast) of multiple directions (every 30°, between 0-330°) were shown a minimum of 10 trials for each orientation and used to determine orientation tuning. Receptive fields were mapped similarly to two-photon calcium imaging experiments without holographic optogenetics (see above).

All visual stimuli were created and presented using PsychtoolBox v3.0 ^70^. Tuning curves were calculated during the experiment, aided by live2p (https://github.com/willyh101/live2p), a custom implementation of CaImAn OnACID (v1.8.8) to perform rigid motion correction and seeded source extraction. Preferred orientation was determined as the orientation corresponding to the maximum mean response to drifting gratings. Visually responsive cells with a high orientation selectivity (>0.5, where OSI = (R_preferred_ - R_orthogonal_) - (R_preferred_ + R_orthogonal_) and R is the mean ΔF/F response) were selected for holographic stimulation.

### Data Analysis

#### Analysis of spiking data

Spiking activity was extracted by filtering the raw signal between 800 and 7000 Hz. Spike detection was performed using the UltraMega Sort package ^74^. Detected spike waveforms were sorted using the MClust package (http://redishlab.neuroscience.umn.edu/MClust/MClust.html). Waveforms were first clustered automatically using KlustaKwik and then manually corrected to meet criteria for further analysis. With the exception of a few burst firing units, included units had no more than 1.5% of their individual waveforms violating a refractory period of 2 ms.

Individual units were classified as either fast-spiking or regular spiking using a k-means cluster analysis of spike waveform components. Since the best separation criterion was the trough-to-peak latency of the large negative going deflection and clustering is non-deterministic, we defined all units with latencies shorter than 0.36 ms as fast spiking and all units with latencies larger than 0.38 ms as regular spiking. Cells with intermediate latencies were excluded from further analysis. Firing rates were computed from counting spikes in a 1 second window starting 500 ms after the onset of the visual stimulus, which coincided with the onset of the LED during optogenetic suppression trials. Contextual modulation index (CMI) was calculated as CMI = (R_cross_ - R_iso_) / (R_cross_ + R_iso_), where R is the mean response during visual stimulation.

#### Two-photon calcium imaging analysis

Raw image files were motion corrected and ROIs were extracted using suite2p ^75^. ROI selection was manually curated in suite2p based on morphology and fluorescence statistics. Neuropil for each ROI (or putative “cell”) was multiplied by a correction coefficient (0.7) and subtracted from the raw fluorescence. Cellwise fluorescence was minimum subtracted, and ΔF/F was calculated as: ΔF/F = (F-F_0_)/F_0_, where F is the neuropil-subtracted, raw fluorescence signal, and F_0_ is the baseline fluorescence, defined as the 10th percentile of fluorescence within a rolling 1-minute window, applied continuously across the time series, for each cell. For a subset of experiments **(Figures 2 and S2-S4)**, calcium traces were deconvolved using OASIS with a L1 sparsity penalty ^43,76^ and reported as event rates.

Retinotopic locations for each cell were estimated by calculating the mean responses to each grid location and then fit with a 2D gaussian. Cells were then classified as “aligned” if they were within 10° of the stimulus center in retinotopic space. Tuning curves were generated by calculating the mean response to each orientation.

Preferred orientation was determined as the orientation corresponding to the maximum mean response of the tuning curve. Orientation selectivity index (OSI) was calculated *post-hoc* using a vector sum approach, OSI = sqrt(∑_i_R_i_cos(2θ_i_)^2^ + ∑_i_R_i_sin(2θ_i_)^2^) / ∑_i_R_i_, where R_i_ are the responses to each orientation, θ_i_ in radians. CMI was calculated similarly to the analysis of spiking data (see Methods above) at the preferred orientation for each cell.

#### Two-photon holographic optogenetics analysis

Putative SST interneurons were manually identified from a mean image collected at 830 nm. For *post-hoc* analysis, holographic targets and BFP-positive SST cells were matched to suite2p extracted cells by the source minimizing the Euclidean distance between centroids. Offline analysis of visual responses were performed similarly to two-photon calcium imaging analysis (see above). Cells were considered visually responsive if they had a significantly different response (p < 0.05, ANOVA) to visual stimuli. Tuning curves for each cell were constructed by collecting the average mean ΔF/F response during visual stimulation across a minimum of 10 trials. Preferred orientation and OSI were calculated as described above. Only cells that were determined to be visually responsive and orientation-tuned (OSI > 0.33) were considered in subsequent analysis. Targeted cells and cells located closer than 15 µm radially from a targeted cell on any z-plane were also excluded from further analysis, based on existing calibrations and characterization of our microscope ^3^. Tuning curves and preferred orientation of SSTs were preferred similarly to PCs, as described above.

#### Statistics

Non-parametric statistical tests were performed, unless indicated otherwise. Individual field of views (FOVs) may come from the same mouse but from a different area within V1 or z-plane and therefore consist of different neurons. No statistical test was performed to predetermine sample sizes. Targeted cells passing selecting criteria were randomly selected as targeted neurons and blind to the experimenter during data collection. Batch analysis scripts were performed *post-hoc* across experimental conditions, effectively blinding the experimenter during data analysis. Source code and data to replicate the figures in the manuscript is available at https://github.com/willyh101/iso-cross. Other source code used in analysis is available at https://github.com/willyh101/analysis. In instances where plot axes are clipped to aid in visualization, statistics were performed on the unclipped and complete dataset.

## Acknowledgments

This work was supported by National Institutes of Health (NIH) grant F32-EY031977 (WDH), F32-EY034022 (MS), NIH grant R01-EY023756 (HA), and U19NS107613 (HA). J.V. was supported by grants from the Swiss National Foundation (P300PA_164719 and P2FRP3_155172) and the DFG (VE 938/2-1, INST 39/1422-1).

This content is solely the responsibility of the authors and does not necessarily represent the views of the National Institutes of Health. We thank Janine Beyer, Karthika Gopakumar, and Savitha Sridharan for technical support.

## Author Contributions

All authors conceived of the study. H.A. and W.H. wrote the manuscript. W.H. performed all two photon holographic optogenetics experiments. H.A. performed in vivo whole cell recordings from pyramidal cells. J.V. performed all extracellular multi-electrode array physiology and one photon optogenetics. D.M. performed all two photon calcium imaging with one-photon optogenetics. M.S. performed all targeted two-photon patch clamp experiments. W.H. prepared all figures and performed all statistical analyses.

---

**Figure S1:**
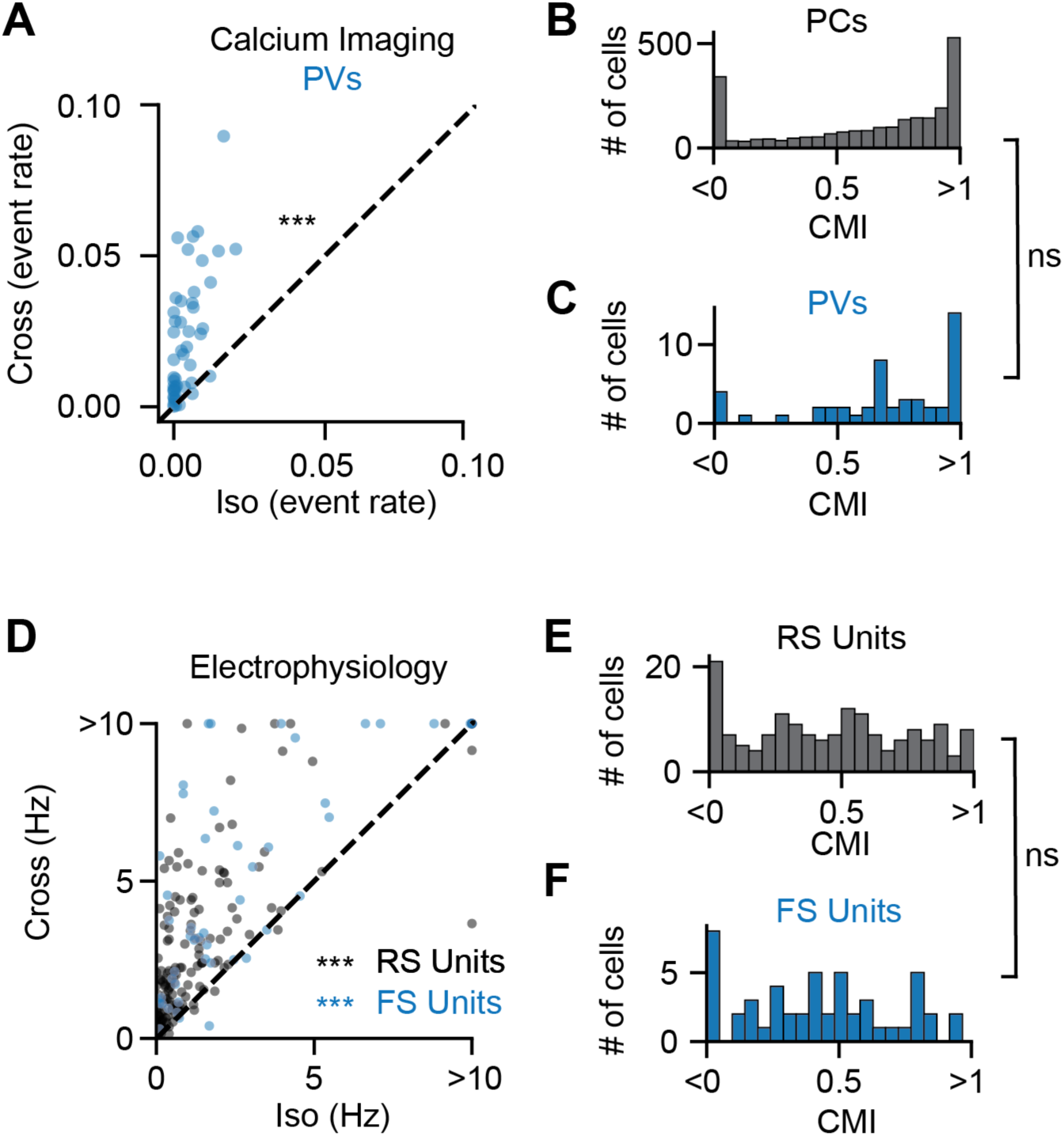
Orientation-tuned surround suppression in both PCs and PVs, and RS and FS units. **A:** Scatter plot of deconvolved calcium responses to cross and iso stimuli taken from PV cells (n=49 cells, p<10^-5^, Wilcoxon signed-rank test). **B:** Distribution of CMI in PCs (n=2,329 cells) taken from *in vivo* calcium recordings. **C:** Similar to **B**, except for PVs (n=49 cells). p=0.168, Mann-Whitney U-test **D:** Scatter plot of cross and iso responses from extracellular electrode recordings to cross and iso stimuli for both RS (black, n=158 units, p<10^-5^, Wilcoxon signed-rank test) and FS units (blue, n=51 units, p<10^-5^, Wilcoxon signed-rank test). **E:** Distribution of CMI in RS units (n=158 units). **F:** Similar to **E**, except for FS units (n=51 units). p = 0.430, Mann-Whitney U-test. ***p<0.001; ns, not significant

**Figure S2:**
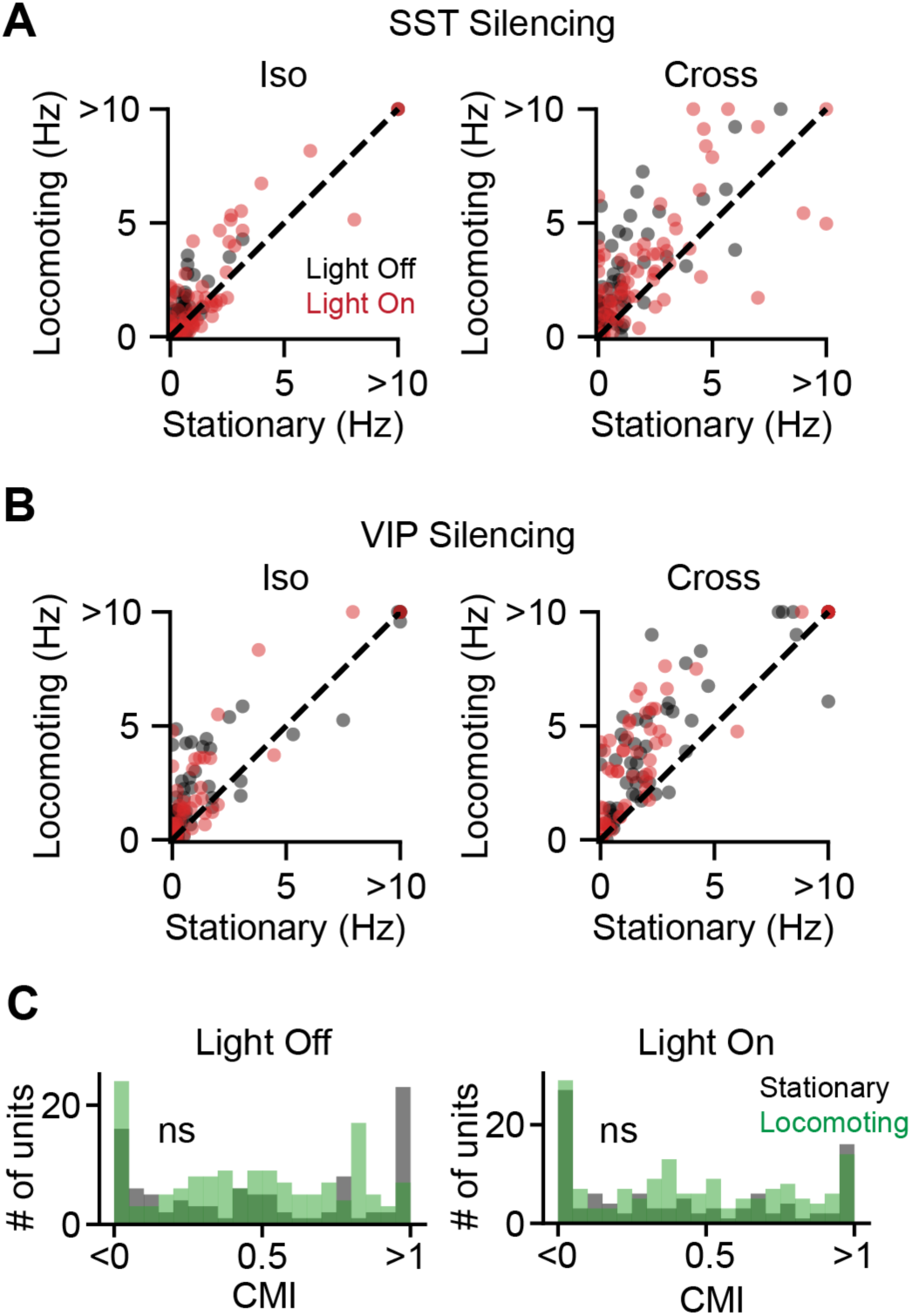
Comparison of firing rates and CMI between stationary and locomoting trials. **A:** Scatter plots comparing RS unit firing rates during SST silencing during iso (left) and cross (right) visual stimuli during locomoting and stationary states. Black points represent responses during control trials (light off) and red points represent firing rates during SST silencing (light on). **B:** Similar to **A**, except for RS units during VIP silencing. **C:** Distribution of RS-unit CMI during stationary (black) and locomoting (green) states for both light off (left) and light on (right) conditions (CMI, light off: stationary, n=98 RS-units; locomoting, n=150 RS units, p=0.393, Mann-Whitney U-test; CMI, light on: stationary, n=101 RS-units; locomoting, n=154 RS-units, p=0.350, Mann-Whitney U-test). ns, not significant

**Figure S3:**
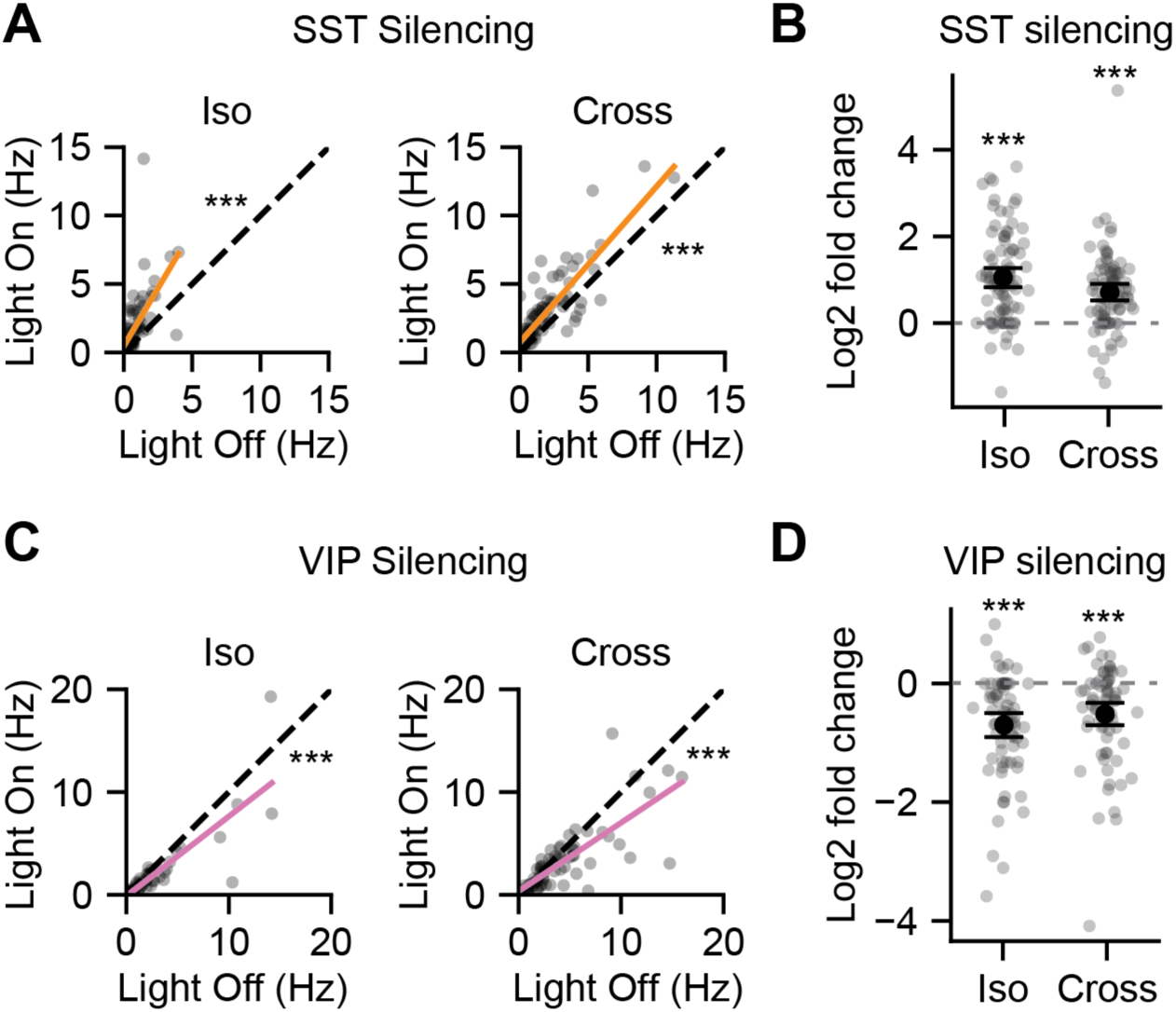
Effects of VIP and SST silencing on putative pyramidal cell firing rates. **A:** Scatter plot depicting the effect of SST silencing on RS unit firing rates (black dots, n=79 RS units) in both the iso (left, p<10^-5^, Wilcoxon signed-rank test) and cross (right, p<10^-5^, Wilcoxon signed-rank test) conditions. Orange line depicts the linear regression between the light on and light off conditions, computed using the Moore-Penrose pseudoinverse. **B:** Quantification of change in RS unit firing rates (n=79 RS units) in response to SST cell silencing to both iso (one-sample Wilcoxon signed-rank test, p<10^-5^) and cross (one-sample Wilcoxon signed-rank test, p<10^-5^) conditions. **C:** Scatter plot depicting the effect of VIP silencing on RS unit firing rates (black dots, n=70 RS units) in both the iso (left, p<10^-5^, Wilcoxon signed-rank test) and cross (right, p<10^-5^, Wilcoxon signed-rank test) conditions. Pink line depicts the linear regression between the light on (y-axis) and light off (x-axis) conditions, computed using the Moore-Penrose pseudoinverse. **D:** Quantification of change in RS unit firing rates (n=70 RS units) in response to VIP cell silencing to both iso (p<10^-5^, one-sample Wilcoxon signed-rank test) and cross (p<10^-5^, one-sample Wilcoxon signed-rank test) conditions. ***p<0.001

**Figure S4:**
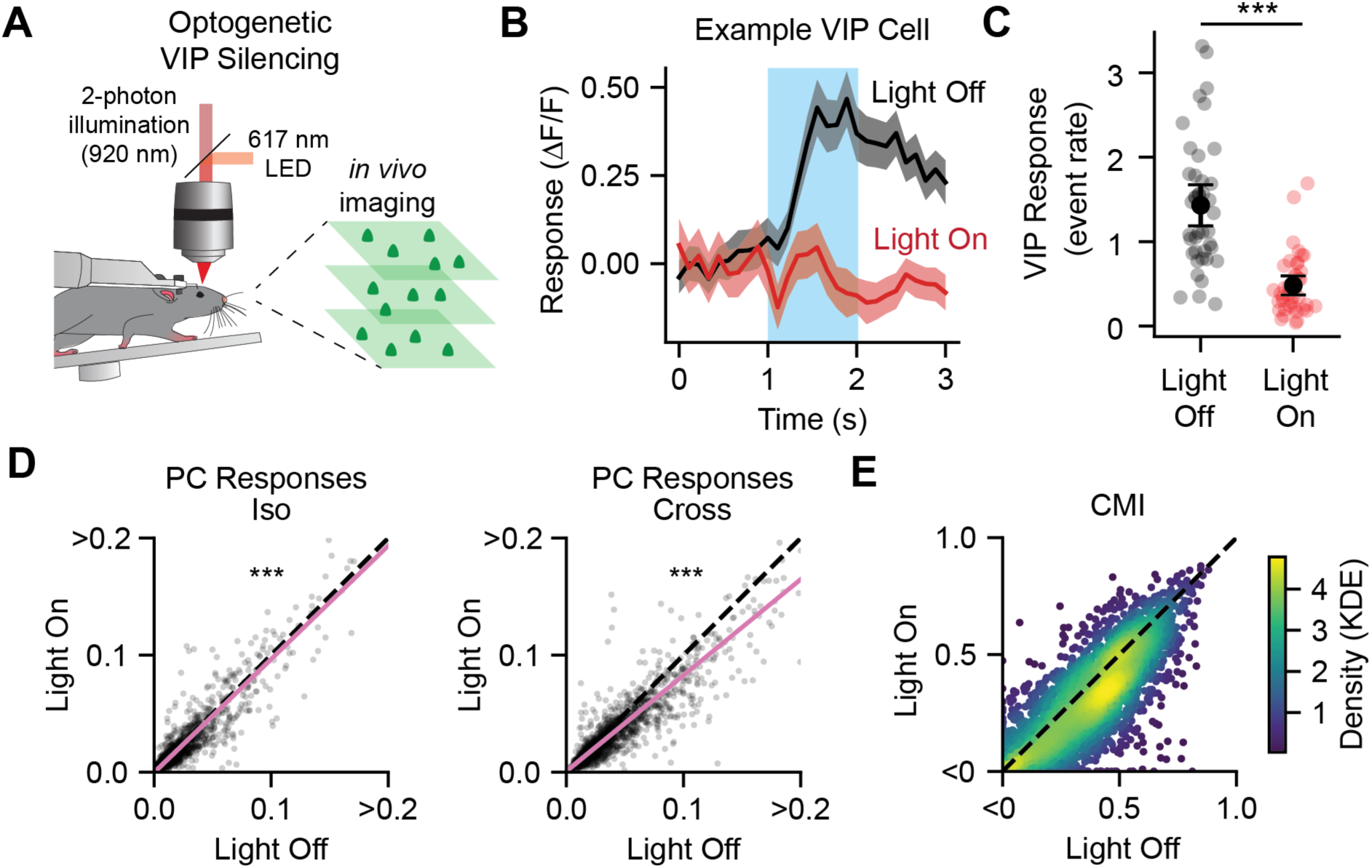
Optogenetic silencing of VIP cells. **A:** Schematic depicting approach for simultaneous *in vivo* 2-photon calcium imaging and one-photon optogenetic VIP silencing. **B:** Visually evoked response (mean ΔF/F) of a representative VIP cell during optogenetic silencing trials (“light on”, red) and control trials (“light off”, black). Light blue shading depicts duration of the visual stimuli. **C:** Quantification of **B** for all retinotopically aligned VIP cells (n=41 cells, p<10^-5^, Wilcoxon signed-rank test). **D:** Scatter plot of visually evoked responses (deconvolved event rate) of PCs (n=2,408 cells) during control (“light off”, x-axis) and optogenetic VIP silencing (“light on”, y-axis) in response to iso (left, p<10^-5^, Wilcoxon signed-rank test) and cross (right, p<10^-5^, Wilcoxon signed-rank test) gratings. Pink lines depict the linear regression between light on and light off conditions, computed using the Moore-Penrose pseudoinverse. Axes have been cropped to a max event rate of 0.2 for clarity. **E:** Scatter plot comparing CMI of PCs in light on (y-axis) and light off (x-axis) conditions (n=2,408 cells, p<10^-5^, Wilcoxon signed-rank test). Individual points are colored by their kernel density estimate (K.D.E.) to highlight regions of higher data concentration. ***p<0.001

